# Initial contact shapes the perception of friction

**DOI:** 10.1101/2021.09.20.461039

**Authors:** Laurence Willemet, Khoubeib Kanzari, Jocelyn Monnoyer, Ingvars Birznieks, Michaël Wiertlewski

## Abstract

Humans efficiently estimate the grip force necessary to lift a variety of objects, including slippery ones. The regulation of grip force starts with the initial contact, and takes into account the surface properties, such as friction. This estimation of the frictional strength has been shown to depend critically on cutaneous information. However, the physical and perceptual mechanism that provides such early tactile information remains elusive. In this study, we developed a friction-modulation apparatus to elucidate the effects of the frictional properties of objects during initial contact. We found a correlation between participants’ conscious perception of friction and radial strain patterns of skin deformation. The results provide insights into the tactile cues made available by contact mechanics to the sensorimotor regulation of grip, as well as to the conscious perception of the frictional properties of an object.

---

We lift glasses of water, regardless of whether they are empty or full and whether they are dry or wet. The sensorimotor mechanisms responsible for this astonishing performance are far from being understood. The grip forces required to lift an object are known to be unconsciously regulated to a value typically 20% above what would cause slippage (1). Remarkably, this regulation starts from the moment our fingers touch the surface. It has been shown that just a hundred milliseconds of contact with a surface are enough to start adjusting fingertip forces to friction. Humans provide larger grasping forces if the surface is made of slippery silk but smaller if it is made of sandpaper since it provides better grasp (2, 3). It has been further demonstrated that it is friction and not texture, which determines these adjustments (3). Since 1 mm of indentation of the fingertip is sufficient to reach 80% of the final gross contact area, and that fingers often move faster than 10 mm/s toward an object, within this time frame the sensorimotor system already should be able to extract some estimates of the frictional properties from the initial deformation of the finger pad, before any net load forces start developing.

On a physical level, the overall so-called *frictional strength* of the contact is given by the number of asperities in intimate contact and their individual shear strength (4–6). It is the measure of the maximum lateral force on the contact that will lead to slippage. This frictional strength is the main determinant in regulating grip force applied to lift an object of a given weight (3). Failure to properly assess the frictional strength of the surface at initial contact –due to the presence of gloves or anesthesia for instance– is followed by larger than usual grip forces, consequently increasing the real area of contact (7–9).

Despite its crucial importance, the mechanical deformation that underpins the encoding of the frictional strength on initial contact remains unclear. It is well known that the timing of the impulses of tactile afferents encodes the information related to force direction (10), local curvature (11), edges (12), shapes (13) and also contains information about the frictional strength (14, 15). One hypothesis suggests that, at the mechanical level, micro-slip events at the finger-object interface induce vibrations of the skin (16, 17). Another hypothesis postulates that the sensation of friction is mediated by a radial pattern of skin strain within the contact area. The magnitude of the strain induces internal stresses, which are 21% smaller on slippery surface than on high-friction surface (18).

Interestingly, roboticists have leveraged these findings to estimate friction on initial contact from the gradient of the lateral traction field. This metric is used to control the force applied by robotic grippers to soft and fragile objects (19–21). In haptic rendering, it is possible to produce tactile sensations by releasing the accumulated stress using ultrasonic friction modulation (22). However, the perception of the frictional strength with a single normal motion is not as salient. Khamis et al. recently showed that participants were unable to differentiate a 73% reduction in friction of a glass plate when it was pressed against their fingertips by a robotic manip-ulator (23).

Friction is consciously perceived in a passive condition only when the plate starts sliding (24–26). The change in the frictional state from stuck to sliding is perceived after a global lateral displacement of 2.3 mm (27). This transition induces large deformations of the skin along with a particular strain pattern (25, 28–30). These results suggest that large or rapid deformations can elicit a tactile sensation, but the quasi-static radial strain pattern is too subtle to induce a reliable percept.

We hypothesize that the frictional strength can be perceived when actively touching the surface. Active exploration is known to promote acute sensitivity (31–33). We present evidence that during the first instant of contact between the finger and an object, a radial strain pattern exists. Its magnitude is affected by the interfacial friction and correlates with the perception of friction. Combined with the results of the motor control literature, a picture emerges explaining the mechanical basis upon which friction is encoded.

## Results

Fourteen participants were asked to actively press down on a glass plate and gauge its frictional resistance. The frictional resistance of the plate against the skin was controlled by ultrasonic lubrication (34), allowing for repeatable stimuli where the surface topography and physico-chemistry remained unchanged.

The apparatus combining a friction plate and an optical system is shown in Fig.1A and construction details are presented in Materials and Methods. The movement of the participants was constrained by a linear guide attached to their finger, preventing any lateral movement (Fig. S2). To validate the reduction in friction, we first asked participants to slide across the plate while the ultrasonic lubrication was modulated (see Fig. S1A). When the finger was steadily sliding and the amplitude of the ultrasonic wave was changed from *α* = 10^−3^ μm to *α* = 3 μm, the coefficient of friction varied from *μ* = 0.81 down to *μ* = 0.18, leading to a 78% relative reduction in friction (Fig.1B).

**Fig. 1.**
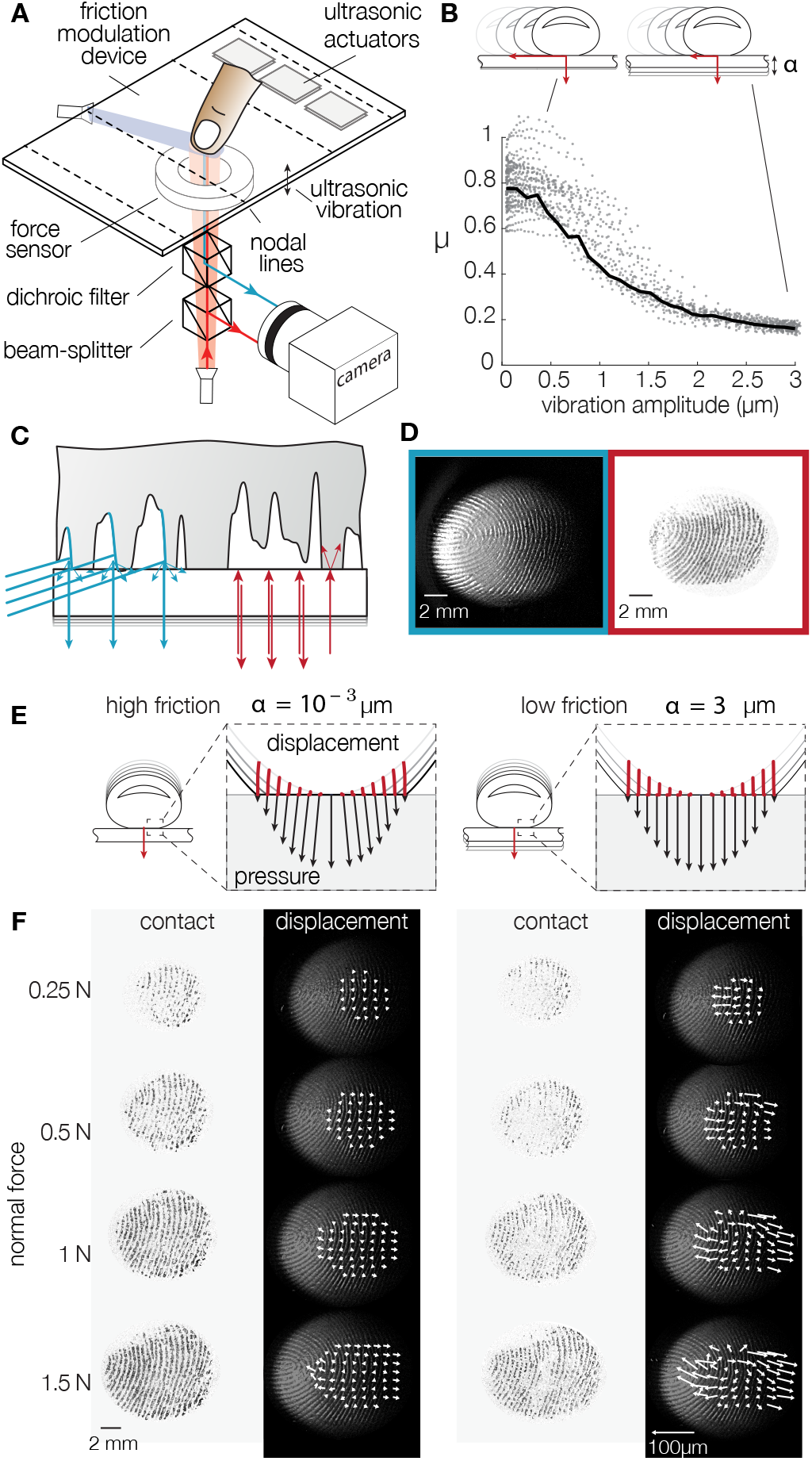
**A**. Experimental setup. The friction between the fingertip and glass plate is reduced in the presence of flexural ultrasonic waves. A dual-illumination setup where blue light illuminates the skin at a 20°angle, and red light is normally incident to the glass surface. **B**. When a fingertip slides across the glass plate, the friction coefficient is reduced with increasing ultrasonic amplitude. The black line represents the median friction coefficient. **C**. Close-up view of the illumination combining a dark-field blue light to highlight the fingerprint ridges, and a red light, coaxially oriented with the camera, to illuminate only the asperities of the skin in intimate contact with the glass plate. **D**. Typical images of the fingertip profile (left) and the asperities in intimate contact (right). **E**. Presumed deformation of the skin when pressed against the surface in high- and low-friction conditions. Points trajectories are shown in red. The black arrows represent the pressure and traction exerted by each point on the surface. **F**. Images of the intimate contact and skin deformation for increasing normal forces (top to bottom) and the highest (left) and lowest (right) friction. The white arrows show the displacement of reference points, scaled up tenfold.

### Empirical skin deformation

In order to accurately measure the plate-fingertip interaction, we used a bespoke illumination apparatus that highlights the topography of the skin, while synchronously showing the micro-junctions that comprise the real area of contact. While participants were pressing down, the motion of individual points on the surface of the skin was tracked using the images from the blue grazing illumination. The high-contrast images created with the coaxial red illumination show the micro-junctions formed by the contact at the interface, providing a temporal reference of the instant when a particular point was in intimate contact. The interaction of the light sources at the skin-plate interface is illustrated in Fig.1C, and the resulting images are shown in Fig.1D.

We can estimate the expected strain with a simple geometrical model, described in Fig. S4. Upon compression against a flat surface, the skin of the fingertip changes from a quasi-hemisphere to a flat disk. It ensues a volume reduction, which can build-up stress at the interface, if friction is high. Conversely, if the surface is slippery, a noticeable deformation is observed (see Fig.1E). This model estimates a 10% compressive strain when the fingertip is indented by 3 mm.

The mechanical behavior of the finger observed during the experiment is qualitatively consistent with the prediction of the geometrical model. Fig.1F shows the evolution of the *real* area of contact constituted by the micro-junctions and the movement of the skin in a high-friction and a low-friction condition for a typical trial. Notably, the real area of contact, shown against a white background, grows with increasing normal force, and its brightness depends significantly on the level of friction reduction. This observation is consistent with previous works and with the adhesive theory of friction, in which the sliding friction force is a function of the real area of contact made by all the individual asperities in intimate contact (34, 35). The displacement vector fields 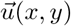 are computed from the difference in position between the final image and the moment when a particular point is detected to make contact. For typical trials a noticeable difference in skin movement between the high- and the low-friction conditions is found, see Fig.1F and movie M1.

### Friction discrimination performance

Participants were asked to compare the slipperiness of the same surface presented with different levels of friction. The friction of the plate was set by the amplitude of ultrasonic vibrations. The reference stimulus was the highest friction when the plate vibrates with an 10^−3^ μm amplitude. The comparison stimuli covered the range of amplitudes from 0.5 to 3 μm at intervals of 0.5 μm, with each stimulus appearing 10 times. The reference and the comparison were presented in random order. After pressing twice on the surface, participants had to indicate which stimulus they felt was the most slippery, following a typical 2-alternative forced-choice protocol. The procedure is depicted in Fig.2A, and movie M2 shows a typical trial.

**Fig. 2.**
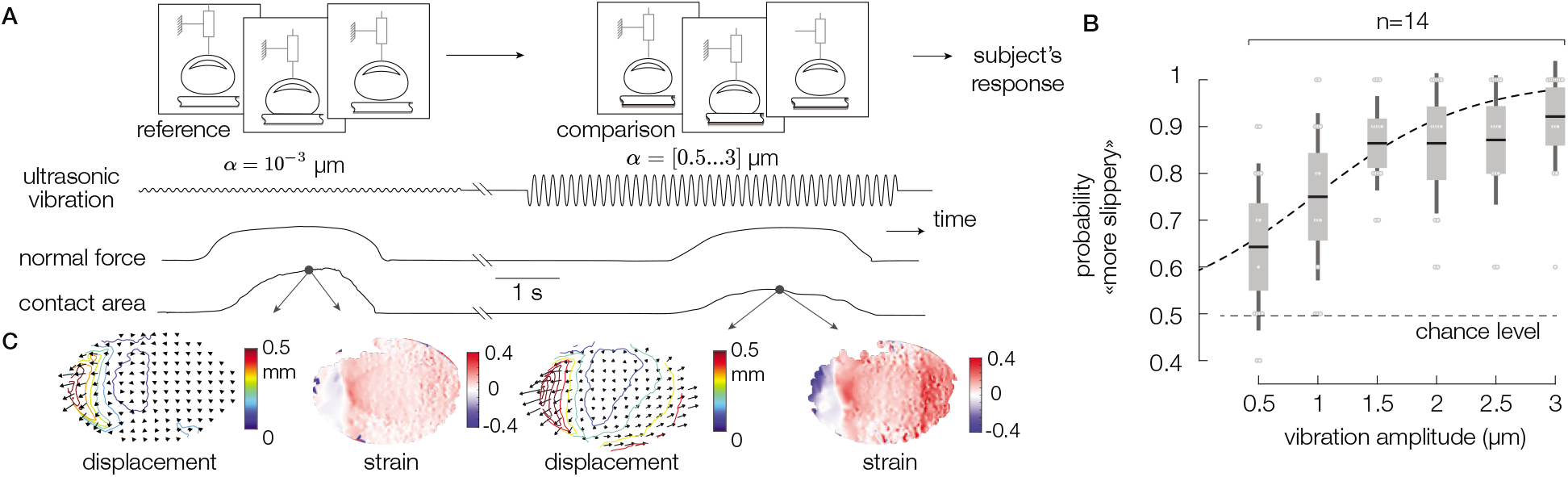
**A**. Experimental protocol. Participants were asked to compare the slipperiness of two surfaces. The reference and comparison are presented in random order. **B**. Probability of a participant perceiving the difference in friction as a function of the amplitude of vibration of the ultrasonic lubrication. The higher the amplitude of ultrasonic vibration is, the lower the friction coefficient. The dashed line represents the fit with a psychometric function. Individual performances are represented as grey dots. The chance level is represented by the dashed line. **C**. Displacement and resulting strain of the skin for the cases of high friction (left) and low friction (right).

We computed the probabilities of responding that the comparison stimulus was the most slippery, and the mean friction discrimination performance for all subjects is reported in Fig.2B. Despite the considerable inter-subject variability, there is a significant effect of plate vibration amplitude on the mean success rate (Repeated-measures ANOVA, F(5,55) = 4.77, p = 0.0011). The results were fitted with a psychometric function, from which we extracted the 75% detection threshold. Participants were able to discriminate the difference of friction, with differences in vibration amplitude as low as 1.13 ± 0.69 μm, which corresponds to a reduction of the real contact area of only 8%.

### Contact and friction modulation

The contact between the finger and the glass plate initially started towards the center and expanded radially, for all trials. In the low-friction condition, the center of the contact experienced ultrasonic levitation, creating areas where skin asperities were not in intimate contact with the plate. The real area of contact, which measures the amount of contact that contributes to frictional strength (34), was reduced by 38% for the maximal vibration amplitude of 3 μm (Fig. S1E). As the number of asperities in contact decreased, the skin could freely expand in the lateral direction, see Fig.2C.

### Friction influences skin deformation

The contact area and displacement field in both high- and low-friction conditions are shown in Fig.3A. In the low-friction case, the regions where the contact was virtually non-existent matched the locations of the regions of maximal displacement of the tracked points. The amount of contact area was measured via the local brightness of a 10-pixel radius circle around each of the tracked points. The displacement of each point positively correlates with the local brightness, hence with the local density of asperities in intimate contact, see Fig.3B (Spearman’s coefficient of 0.58). This relation provides evidence that at the scale of fingertip features, friction does influence the lateral mobility.

**Fig. 3.**
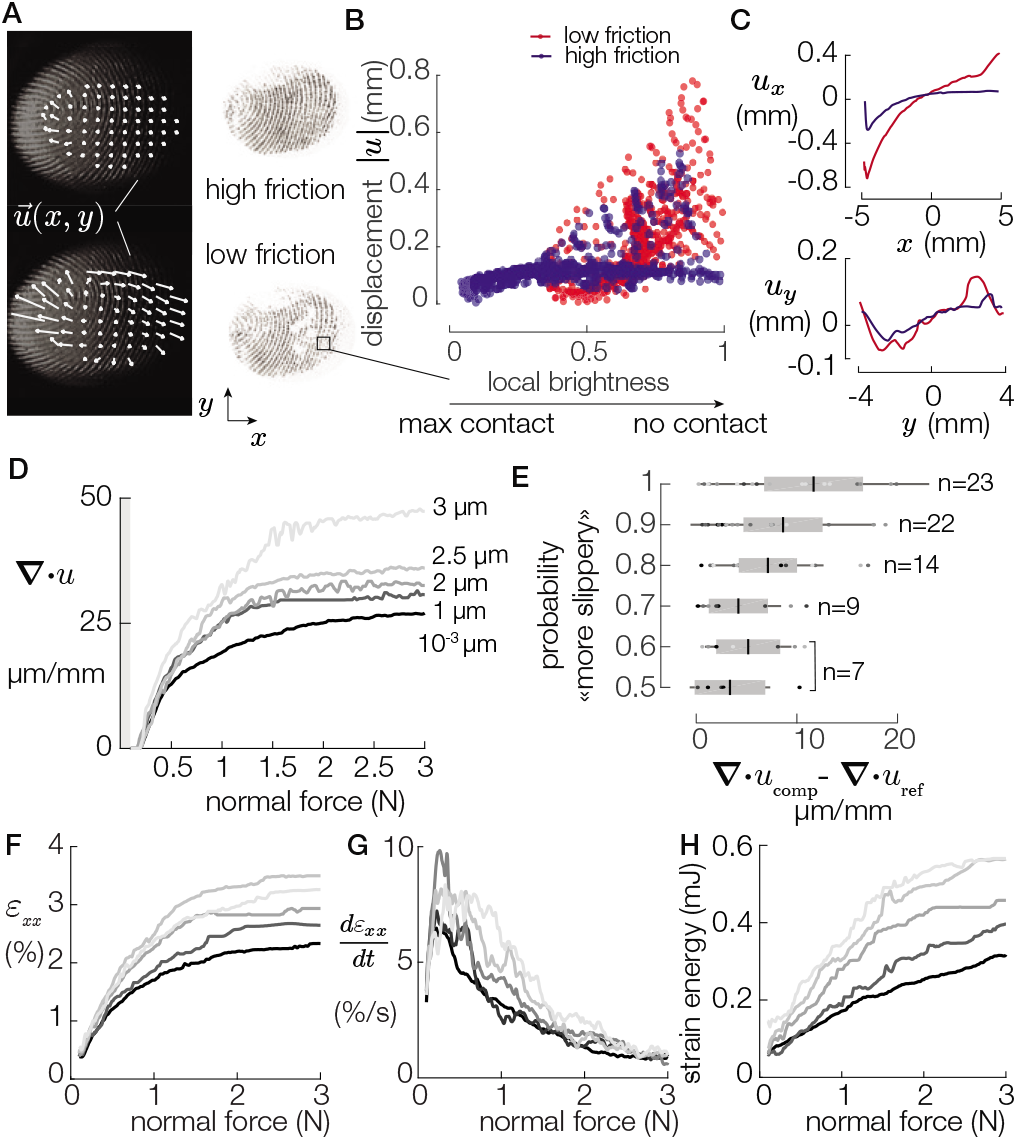
**A**. Typical images of the high- and low-friction trials. **B**. Displacements of a grid of points plotted against the local brightness. Displacements and local brightness are positively correlated (Spearman’s coefficient = 0.5761, *P* < 0.0001). **C**. Displacements of this typical grid of points along the x- and y-axes. **D**. Median divergence for each vibration amplitude. **E**. The probability to answer comparison is the most slippery are plotted against the median divergence differences. Darker colors represent the smaller vibration amplitudes. **F**. Median longitudinal strain for each vibration amplitude. **G**. The strain-rate peaks after 0.4 N for each vibration amplitude. **H**. Evolution of the strain energy for various coefficient of friction.

The lateral displacements of the skin along the *x* and *y* axes are shown in Fig.3C for the low- and high-friction cases. The projection along the central axis reveals that the center of the contact experiences a deformation gradient whose value depends on the frictional state. To explore the effect of friction on the displacement field, we decomposed it into a constant field, a divergent field and a rotational field (see Fig. S6 and S13). The mean divergence of these micro-displacements was computed such that:

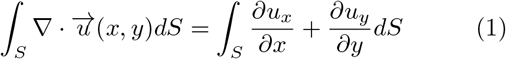

where *u_x_* and *u_y_* are the x and y components of the displacement vector *u*(*x*, *y*) respectively, and *S* is the apparent area of contact. Intuitively, the averaged divergence of a vector field captures its outward or inward flux. A positive divergence implies that the finger expands radially.

Fig.3D plots the median across trials of the average divergence for increasing plate vibration amplitude. The divergence grows with the normal force. The rate of growth is positively correlated with the vibration amplitude (Spearman’s coefficient = 0.115, p < 0.0001). The growth of the average divergence is notable at the early stage of fingertip compression and hits an inflection point after 1 N. After this inflection point, the dependence on friction is more pronounced. Above 2 N, the curves flatten, likely due to saturation of the compres-sion of the fingertip pulp (36, 37). Despite the saturation above 2 N, the differences in average divergence are significant (ANOVA, F(6,1569) =4.85, p = 10^−5^), with values twice as large for the low-friction case (3 μm) than for the high-friction case (10^−3^μm). Large divergence reflects that the skin moves significantly without friction. In the high-friction case, the low divergence values signal the presence of residual radially distributed stress of the skin.

### Skin deformation and friction perception

The global lateral displacement of the skin, computed from the median of the vector field at each time instant, and the peak force vector angle did not significantly influence the response of the participant (Spearman’s correlation, *p* = 0.2 and *p* = 0.03 respectively), see Fig. S7C,D. Because every participant was free to actively press against the surface, the recorded normal forces (avg = 5.5 ±3.5 N) and the duration (avg = 1.47 ±0.39 s) showed significant variations. We found no significant relationship between the friction discrimination and normal force or the duration of contact (see Fig. S5). However, we found a significant influence of the force rate on the participants’ answers for the vibration amplitude *α* <= 2 μm (Linear Mixed Model, p = 0.018) (Fig. S5I). The kinematics of the exploratory procedure play a significant role for the low vibration amplitudes.

Nonetheless, an ideal observer analysis shows that the average divergence of the skin deformation was the most salient predictors of participants’ responses (see Supplementary Materials). Fig.3E shows the difference in divergence between the reference stimulus and the comparison stimulus as a function of participants’ discrimination performance. The probability of correctly identifying the most slippery stimulus was positively correlated with the amount of diverging skin deformation observed (Spearman’s coefficient = 0.28, *p* = 0.009). While the correlation is weak, friction was unambiguously discriminated when the skin experienced the largest inter-stimulus difference in divergence.

### Strain energy and mechanoreceptors thresholds

It is worth considering whether the amount of skin deforma-tion is enough to induce a supraliminal response. The finger asperity which experienced the maximal displacement moves with a speed of 1.93 ±2.8 mm/s (mean ±std), see Fig. S5K.

We estimate the stimulation of the mechanoreceptors by computing the strain components, according to the method described in Supplementary Materials. The data show that the participants’ skin is subjected to a longitudinal strain whose magnitude depends on the vibration amplitude (Spearman’s coefficient ρ = 0.17, p < 0.0001), see Fig.3F. The strain magnitude estimates fall between 2 and 4%, which is sufficient to change firing rate in FA and SA afferents (38). Similarly the dynamics of the stimulation shows significant differences between friction condition, see Fig.3G. The strain-rate peaks at 8.5 ±1.7%/s when the normal force reaches 0.37 ±0.7 N for all friction conditions. This is compatible with the evidence in literature that a stimulation with a strain rate higher than 8%/s elicits a response in all afferent types (39).

### Predictions from a mechanical model

We developed an axisymmetric spring-damper model illustrated in Fig.4A, to estimate the stress experienced by the skin. The model captures the large deformation of the skin, its viscoelastic behavior using Kelvin-Voigt material and the local elastoplastic frictional interaction at the interface. This model is detailed in Supplementary Materials.

**Fig. 4.**
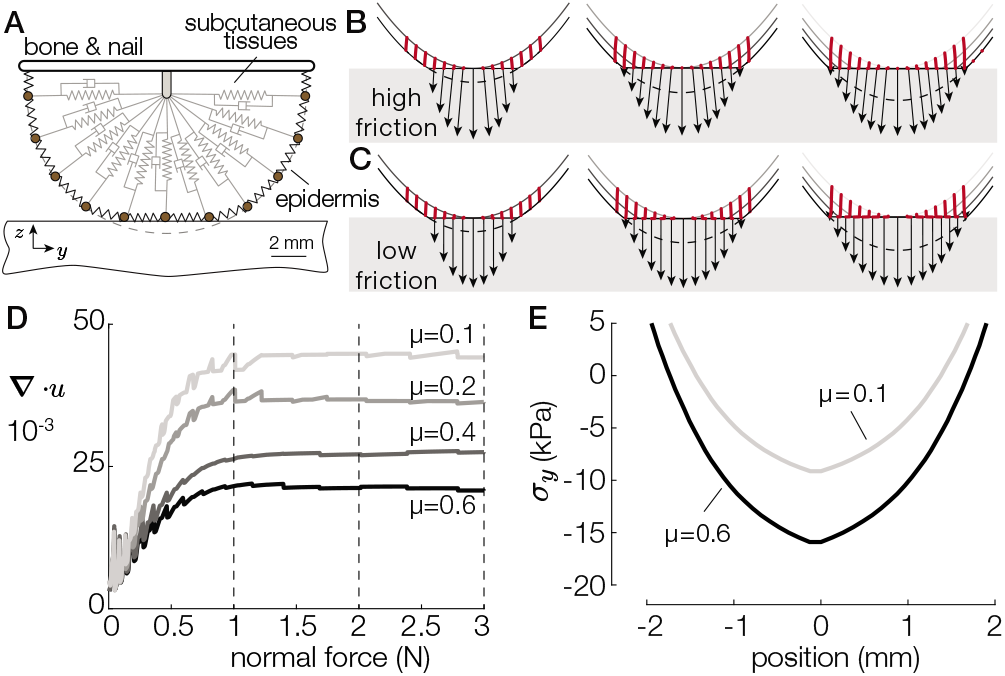
**A**. The finite difference model of the fingertip. Surface deformation profiles when in contact with a low- **B** and a high- **C** friction surface for 1, 2 and 3 N. Points trajectories are shown in red. The black arrows represent the pressure and traction exerted by each point when brought in contact with the surface. **D**. Total divergence of the contacted skin as a function of the normal force for friction coefficients varying from 0.1 to 0.6. **E**. Stress profiles at 3 N of normal force for a high- and a low-friction condition.

We simulated the interaction with a flat surface under four different coefficients of friction from 0.1 to 0.6, with a normal force of 3 N applied to the bone element. The simulation was initiated before contact and ran until it reached static equilibrium. M3 shows the results of the simulation for the high- and low-friction conditions. The simulated displacements and interfacial pressure in response to an external normal force of 1, 2 and 3 N for *μ* = 0.1 and *μ* = 0.6 are shown Fig.4B and 4C respectively. Fig.4D shows the simulated divergence of skin displacement for all friction coefficients. The divergence varies from ∇*u* = 0.02 for a coefficient of friction *μ* = 0.6 to ∇*u* = 0.04 for *μ* = 0.1. The model predicts a trend quantitatively similar to the experimental data.

These observed lateral displacements can be explained by the stress acting on each element, caused by friction. The normal component of the interfacial pressure remains identical across frictional conditions. However, the lateral component directly depends on the friction, with the high-friction case seeing 40% larger tangential stress (Fig.4E). The maximum of the stress is located on the center of the contact area and is consistent with the traction observed in (18, 21). In the low-friction case, the low tangential traction results in a free lateral displacement of the skin as the fingertip flattens in contact with the plate. In this case, every point moves outward such that the contact length approaches the initial curved length of the fingertip (dashed line). Conversely, in the high-friction case, the tangential traction constrains the motion of skin in contact. The elements are secured in place once they touch the plate, resulting in little displacement and a 40% increase in stored elastic stress compared with the low-friction case.

## Discussion and conclusion

A short one-second haptic normal force contact was sufficient to allow participants to discriminate the frictional strength of a surface. The results demonstrate that no gross lateral motion of the whole contact area was necessary to elicit the perception of friction. The observers fundamentally relied on cutaneous cues, involving a particular spatio-temporal pattern following an outward expansion quantified by the divergence of the skin deformation.

The results indicate that a causal relationship exists between the lateral deformation during compression and the observed real area of contact. Under low-friction conditions, fewer asperities are in intimate contact, and therefore, they cannot hold the lateral force, inducing local slippage. Conversely, in high-friction conditions, the asperities make sufficiently large contact and thus restrain lateral relaxation of the skin, which causes an accumulation of the elastic stress. These friction-dependent skin deformation changes can be described by a biomechanical model. Psychophysics experiments demonstrated that the magnitude of friction reduction effect correlates with likelihood of subjects identifying the most slippery surface. As there were no net lateral forces present, the pattern of outward skin expansion characterized by divergence was the decisive factor to assess friction when other cues are not available.

We estimate that the action of pressing down against a high-friction surface stores approximately 0.3 millijoules of potential elastic energy in the skin (Fig.3H). This amount is 10 times lower than what was found when detecting slippage during relative motion, where the strain can reach 25%. This result suggests that information about the frictional strength is available well before slippage is detected (27, 28).

The amount of lateral skin deformation during pressing is sufficient to trigger a significant difference in the activation of all types of tactile afferent (15, 40, 41). In this study, the relative speed between the skin and the glass plate at the periphery of the contact is larger than 10 mm/s. This speed, combined with the spatial nature of the deformation pattern, suggests that fast-adapting afferents predominantly contribute to the encoding of friction upon initial contact.

An early estimation of the frictional strength has been associated with an early adjustment of the grip force during precision grasping tasks. New evidence obtained in the current study extends these findings showing that in a well-controlled perceptual task, abolishing all additional contributing factors like lateral force and texture cues, friction discrimination was possible perceptually. This indicates that information about the initial skin deformation pattern can be sufficient to obtain frictional information. However, during object manipulation beyond initial touch, when load forces develop, more sensory signals become available, improving force coordination and making overall adjustments to friction more accurate (15, 18). Gloves and other mechanical filters are well known to affect the regulation of grip forces, resulting in an overcompensation of the safety margin increases regardless of the friction of the surface (8, 14, 18). The presence of this mechanical filters might remove the ability to gauge the divergence of the field during the first instant of contact, hence defaulting motor control to a more robust grasping state.

The skin deformation increases with the applied normal force, and its rate of increase is a function of the friction of the surface. Despite growing at different rates, the divergence of the displacement field reaches a plateau at 2 N of normal force for all friction conditions, which is similar to the level of grasping force at which friction starts to influence the rate of grip force increase (2). This result suggests that during the first instant of contact, grasp control may rely on the measured divergence of the skin deformation.

Interestingly, the perception of the softness of an object during active touch is correlated with the rate of growth of the contact surface (42). Since friction influences the rate of change in the elastic energy, we can conjecture that cross-coupling might exist between softness and friction, with slippery surfaces appearing more compliant to the touch.

Despite having similar levels of friction variations and observed skin displacement up to 0.2 mm in magnitude, previous studies in which participants passively perceived the stimuli showed that the discrimination of friction is a challenging task (23). In contrast, the active exploration procedure of this study, even if constrained, resulted in a fundamentally more successful discrimination of the frictional conditions. The stark difference could be explained using predictive coding theory (43). To determine friction, the observer has to assess the total deformation separating at least these two components, one encoding the indentation magnitude and another related to the lateral deformation encoding the frictional strength. In the active case, observers possess an efferent copy based on which they could predict the dynamics of gross deformation of the fingertip. The ability to predict sensory consequences of own actions (reafference) would enable the nervous system to better extract and isolate sensory signal features related specifically to the diverging deformation pattern, and thus focus attention to frictional cues.

Alternatively, it is possible that a difference in indentation speed may have played the major role determining detectability of the frictional differences. Khamis et al. (23) report a force rate of 1.7 ±0.3 N/s, whereas in this study the force rate is 3.6 ±3 N/s, which would provide a more potent activation of fast adapting afferents (Fig. S5I).

This study establishes the link between skin deformation and performance in a friction discrimination task. Similar to the suggestion in (28), an artificial tactile stimulation stretching the skin radially while the user is pressing down, could indicate the amount of friction. These cues could facilitate the manual control of teleoperated devices or render a virtual sensation of slipperiness. The biomechanics can also inspire the control of robotic grippers and prostheses based on radial lateral skin stretch (19).

## Materials and Methods

### Ultrasonic lubrication

The friction reduction device uses a flexural standing wave to induce micrometric levitation of the skin of the fingertip. The device is composed of a rectangular glass plate vibrating at a frequency of 29.194 kHz in the 1×0 mode, with dimensions of 67 × 50 × 5 mm^3^. The plate is mounted onto an aluminium frame attached to a 6-axis force sensor (ATI Nano 43) to measure forces exerted by the finger with 10 mN accuracy.

### Participants and protocol

Fourteen right-handed volunteers (3 females and 11 males), ranging from 19 to 55 years old, participated in the study. They were naive to the purpose of the experiments and had no previous experience with haptic devices. None of them reported having any skin conditions or perceptual deficits. The study was conducted with the approval of Aix-Marseille Université’s ethics committee (201914-11-003), and the participants gave their informed consent prior to the procedure. Participants sat in a chair in darkness and wore noise-cancelling headphones projecting pink noise, blocking any visual or auditory cues. The last phalanx of their left index finger was connected to a vertical linear guide, preventing any lateral movement (Fig. S2A). The approach angle of the finger was maintained at 30°. The entire session was composed of 2 blocks of 20 min, separated by a 10 min break. Participants stated which stimulus was the most slippery, a correlate of the friction coefficient.

### Data analysis

Force data were synchronized to the images, interpolated to match the time vector of the images, and denoised with a zero-lag 50 Hz second-order low-pass filter. The global displacement was computed for each trial by summing all the displacements in the apparent contact area. Trials in which the global displacement exceeded 0.3 mm were removed from the divergence analysis to prevent participants from using this cue. Seventy-nine trials out of 840 were removed.

The denoised and illumination-corrected images were thresholded using Otsu’s method to measure the number of asperities in intimate contact. The contact surface in mm^2^was computed by summing the number of white pixels scaled by pixel resolution in mm/px (see Fig. S3A).

The 700 most salient features of the fingerprint were tracked from the start until the normal force reached 3 N. The displacements of these features are interpolated on a uniformly sampled rectangular grid (Fig. S3B) to compute the divergence. The evolution of the median of the divergence field quantified the observed expansion.

## ACKNOWLEDGMENTS

We would like to thank Alessandro Moscatelli and Vincent Hayward for their precious insights. This work was supported by the ANR (PHASE 16-CE10-0003), 4TU Soft Robotics Program and the Australian Research Council (DP170100064).

