## Supplementary Materials for "Initial contact shapes the perception of friction"

September 20, 2021

Corresponding Author name: Laurence Willemet


#### SI Materials and Methods

##### Friction modulation and contact area

To demonstrate the ability of the plate to reduce friction, participants were asked to slide their finger over the surface while the amplitude of the ultrasonic carrier was modulated with a 4 Hz sinusoid. The evolution of the normal and tangential forces was measured with a custom-built tribometer. The tribometer relied on a rigid elastic structure, which nanometre-scale deformation was measured via a Fabry-Perot interferometer. See (1) for construction details. This high-precision sensor can resolve forces with amplitudes lower than 1 mN.

Participants were asked to keep the normal force steady around 0.5 N on average. The epochs where the finger was moving from the right to the left and the vibration envelope increased were selected. The friction coefficient was computed from the ratio of lateral to normal forces for each separate epoch (Fig.S1A).

Since no frictional forces were present during the normal indentation by the participants during the 2-alternative forced choice procedure, the friction coefficient cannot be computed from the force ratio. Thus, we used the area of contact as a proxy measurement for friction.

The area of contact of skin on glass can be characterized in two ways; the apparent area of contact, which is the macroscopic area due to the gross deformation of the tissues; and the real area of contact, which is made by summing the contribution of the microscopic scale junctions between the asperities of the skin and the glass plate.

The observed contact areas varied significantly across participants with values ranging from 70.1 ± 4.5 $mm^2$ for the apparent contact area and 23.7 ± 4.5 $mm^2$ for the real contact area. The variation is attributed to differences in skin reflectance, humidity, and fingertip size. The contact areas were normalized to the median size of each individual to compare the results across all participants. The apparent contact area is not affected by the ultrasonic levitation (Fig. S1CD), as previously shown by Wiertlewski et al. (2). However, the normalized real contact area evolved almost linearly with the normal force (see Fig.S1E), and the slope of the relationship was negatively correlated with vibration amplitude (Spearman's coefficient = -0.28, $p < 0.0001$). The correlation is illustrated in Fig.S1F, in which the maximal vibration amplitude of 3 µm caused a 38% reduction in the contact area, consistent with ultrasonic lubrication theories (2) and with friction theories (3). It reveals that fewer asperities were in intimate contact, thus potentially allowing more lateral movement of the skin unimpeded by friction.

##### Images acquisition and processing

The mechanical interaction with the participant's skin was visualized with a custom-made optical system (fig. S2B). A 450 nm blue light (Thorlabs M455L3) illuminates the fingerprints at a shallow angle of 20°. A 660 nm red light (Thorlabs M660L4) is shone via a beam-splitter, so its principal axis is orthogonal to the surface of the glass and parallel to the optical axis of the camera. This type of illumination used in (4) leverages the frustration of the 4% reflection of the glass surface by the skin to image the asperities in great details. A dichroic filter (Thorlabs DMLP 550) and a set of mirrors spatially separate the two illumination sources. The images were captured at 300 frames per second by a high-speed camera (Phantom VEO E310) with a resolution of 512x640 pixels covering a total area of 16x21 mm.

Gathering the real contact area followed a three-stage process (Fig. S3A): i) The raw image of the contact area was first normalized to a reference image containing only the illumination function. ii) Once the uniformity of the light was restored, a 2d median filter with a 9x9 kernel removes salt and pepper noise. iii) Otsu's method provides a thresholded image of the border of contact, from which we extracted the real contact area by summing the pixels.

Once the image of the contact was found, we computed the deformation field from the topographic image (Fig S3B). Robust features of interest that lied in the apparent area of contact were tracked. To do so, the image of contact was registered according to the topographic image, using a calibration object containing 3 non-aligned points. The registered image followed the same treatment as the one used to compute the contact area. At the end, the binarized contact image was dilated with a radius of 8 pixels and an ellipse was extracted from this image.

Contrast of the topographic image was adjusted, and the contour was sharpened. The algorithm of Shi & Tomasi (5) was used to select 700 optimal features to track inside the ellipse of contact. Then, these features were tracked using Lucas & Kanade algorithm (6). The tracker tracks each point from the previous to the current frame and computes the bidirectional error, which is the distance in pixels from the original location of the points to the final location after the backward tracking. If the maximal bidirectional error exceeds 1 pixel, the point is considered to be not reliably tracked. The image showing the micro-junctions formed by the contact at the interface provides a temporal reference to mark when the tracked points were in contact. Subtracting the position of each point once it first touches the plate, we obtained the 2-dimensional displacement field. From this vector field, the global displacement of the finger was obtained by computing the median value of the travelled distance by all tracked points. Finally, the divergence field was computed at each time instant and for each point once they were in contact with the plate using equation 1 in the manuscript and the *gradient* function in Matlab.

#### Strain computation

The strain components were obtained via the same procedure as in (7). A Delaunay triangulation was first constructed with the 700 tracked points, only considered once they enter in contact with the plate. This triangulation is illustrated in Fig.S8A and C. Then, we used the following formulas to compute the strain components of each triangle.

$$\begin{aligned}
\epsilon_{xx} &= \frac{\partial u}{\partial x} + 0.5\left[(\frac{\partial u}{\partial x})^2 + (\frac{\partial v}{\partial x})^2\right] \\
\epsilon_{yy} &= \frac{\partial v}{\partial y} + 0.5\left[(\frac{\partial u}{\partial y})^2 + (\frac{\partial v}{\partial y})^2\right] \\
\epsilon_{xy} &= 0.5\left[\frac{\partial u}{\partial y} + \frac{\partial v}{\partial x}\right]
\end{aligned} \tag{1}$$

The strain energy densities $u_d$ were computed for each triangle based on average values of Young's modulus and Poisson's ratio, respectively equal to 1 MPa and 0.4. Note that these values can nonetheless vary from one participant to another.

$$u_d = \frac{E(1-\nu)}{2(1+\nu)(1-2\nu)}\,(\epsilon_{xx}^2 + \epsilon_{yy}^2) + \frac{E\nu}{(1+\nu)(1-2\nu)}\,\epsilon_{xx}\,\epsilon_{yy} + \frac{E}{1+\nu}\,\epsilon_{xy}^2 \tag{2}$$

The total strain energy on the whole contact area was obtained by integrating the strain energy densities on a volume, assuming that the strains are uniform for a given depth of 2 mm (7).

$$U = \int u_d\; dV \approx 0.002 \int u_d\; dS \tag{3}$$

#### Skin deformation model

The model is built to be as parsimonious as possible, while retaining predictive power over the observed behavior. It is composed of a chain of massless elements maintained together by elastic springs. This chain can be assimilated to the external layer of the skin (the epidermis). Its shape is maintained using other elastic springs that connect the massless elements to a virtual bone, analogous to the mechanical behavior of the subcutaneous tissues. The two elements on the outside of the membrane are also attached to the bone and model the effect of the rigid nail. Overall, the model resemble the discrete version of a curved elastic membrane on a spring foundation. The viscosity of the skin is modeled by dampers, connecting each particle to the mass of the system.

Let $F_i$ be the force created by all springs and dampers acting on the particle $i$ and $\mathrm{u}_i$ the displacement of the particle $i$. The internal force on each element $i$ is written as following:

$$F_i = -k_m(\mathrm{u}_{i-1} - 2\mathrm{u}_i + \mathrm{u}_{i+1}) - k_t(\mathrm{u}_i - \mathrm{u}_b) - \zeta\dot{\mathrm{u}}_i \tag{4}$$

where $k_m$ is the stiffness of the external layer of the skin equal to 2.5 kNm, $k_t$ is the stiffness of the subcutaneous tissues equal to 0.13 kNm and $\zeta$ is the damping coefficient equal to 0.1. These values enable to match the observed static and dynamic behavior of human finger. Materials properties including young's modulus of human external layer of the skin have already been measured (8, 9). Stiffness of the internal layer was adjusted to fit the observed relationship between normal pressure and contact area (10). A 0.1 damping coefficient fix the time response to 10 ms (9).

As mass and inertia are neglected, the equation of motion is the following:

$$\mathbf{B}\dot{\text{U}}(t) + \mathbf{K}(\text{U})\,\text{U}(t) + F_{\text{ext}}(t) = 0 \tag{5}$$

where U is the state vector of displacements and $F_{\text{ext}}$ is the external forces vector. **K** and **B** are respectively the matrices of springs and dampers dependencies.

The stiffness matrix **K**, is repopulated at each time-step to take into account the geometric changes. Because it depends on the position of each element, the system of equations is essentially non-linear. The displacement vector U and the impedances are decomposed into a normal and a tangential component. For example, the normal and tangential components of spring stiffness depend on the angle between thee surface and the spring $\alpha$ such that $k_m \sin\alpha$ and $k_m \cos\alpha$, respectively.

Equation (6) is solved in discrete time using 4th order Runge-Kutta iterative method such that:

$$\begin{gathered}\mathbf{B}\left(\frac{\text{U}(t+dt) - \text{U}(t)}{dt}\right) + \mathbf{K}(\text{U})\,\text{U}(t) - F_{\text{ext}}(t) = 0 \\ \text{U}(t+dt) = \text{U}(t) + dt\left(-\mathbf{B}^{-1}\mathbf{K}(\text{U})\,\text{U}(t) - \mathbf{B}^{-1}F_{\text{ext}}(t)\right)\end{gathered} \tag{6}$$

External normal forces due to contact were updated using the penalty method. When the element was subjected to a normal force, friction force were computed using Dahl's model (11) such that:

$$s\frac{dF(x)}{dt} = \frac{dF(x)}{dx}\frac{dx}{dt} = \sigma_0 \left|1 - \frac{F}{F_c}\text{sign}(\dot{x})\right|^n \text{sign}(1 - \frac{F}{F_c}\text{sign}(\dot{x}))\dot{x} \tag{7}$$

where $F(x)$ is the friction force function, $F_c$ is the coulomb friction force and $\sigma_0$ is the rest stiffness at equilibrium point $F = 0$, taken equal to 500 here. $n$ is a coefficient that codes how ductile or brittle the material is, equal to 0.7 here. Then $F(x)$ approaches the coulomb friction force $F_c$ as long as $\dot{x} > 0$ and $-F_c$ when the direction of motion is reversed.

The number of elements along the chain is fixed according to the Courant-Friedrich-Lewy (CFL) condition, ensuring the convergence of the system:

$$\frac{v\Delta t}{\Delta x} \leq C_{max} = 1 \tag{8}$$

where $v$ is the maximum magnitude of the velocity. Since the maximal speed approaches 440 m/s and the temporal discretization $\Delta t$ is defined equal to 2.5 µs, the spatial step $\Delta x$ should be higher than 1.1 mm. Taking 501 elements thus ensure the convergence.

### SI Results and Discussion

#### Intuitive explanation of the formation of radial strain

We can build an intuitive understanding why the skin experiences a radial lateral stretch by considering that the fingertip is geometrically approximated to a deformable half-sphere. When this half-sphere enters in contact with a surface, the deformable structure flattens at the interface (fig.S4A). If the friction is considered to be infinitely high, the elements in contact are locked in place and are not able to move laterally. Thus, the length of the arc of the skin $L$ is compressed to fit within the contact area $a$. Both of these dimensions can be estimated from the finger radius $R$ and the normal indentation $\delta$, which depends on how much the finger is pressing on the surface ((9)). Fig.S4B plots both lengths as a function of $\delta$.

$$\begin{gathered}L = R\arccos\left(\frac{R-\delta}{R}\right) \\ a^2 = (R^2 - (R-\delta)^2)\end{gathered} \tag{9}$$

Skin strain $\epsilon$ can be computed with (10). For a 3 mm normal indentation, the skin experiences a 10% lateral compression (fig.S4C).

$$\epsilon = \frac{L - a}{L} \tag{10}$$

#### Influence of the kinematics of the exploratory procedure

Participants were free to press at any normal force and as long as desired. Consequently, the range of normal forces developed by the subjects varied from 1 to 18 N with a mean around 5.5 N (Fig.S5B). The time to reach the peak normal force follows a normal distribution with a mean of 1.47 s and a standard deviation of 0.39 s (Fig.S5C). The total duration of every trial varies from 1 s to 2.5 s (Fig.S5D). In any case, the amount of force applied or the duration of the trial were not significantly correlated with participants' answer (ANOVA, p = 0.31 for normal force, p = 0.99 for time to max force and p = 0.91 for duration). Therefore, these metrics did not give any cues to discriminate friction (Fig.S5E, F and G). However, large normal forces were found to be associated with low probabilities. Our hypothesis is that when participants haven't any valuable cues to discriminate friction, they press harder to induce larger skin deformation.

The force rates applied by the participants follow a normal distribution of mean 3.6 N/s and standard deviation 3 N/s (Fig.S5H). In the bar plot in Fig.S5I, the probability that participants will identify the comparison stimulus as most slippery is shown as a function of the force rate for each of vibration amplitudes. We found that the force rate has a significant influence on the participants' answers for the vibration amplitude $\alpha <= 2$ µm (Linear Mixed Model, p = 0.018). The faster the indentation speed, the more the chance to detect correctly the most slippery stimulus.

#### Influence of global displacement and force vector angle

Lateral global displacements were estimated by computing the median of all vectors in the apparent contact area at each time instant, they represent the constant part of the deformation field, as shown in figure S6. Global displacement takes relatively small values (avg=0.08 mm ± 0.10 mm SD) (Fig.S7A). In addition, the global displacement of participants' fingers cannot be considered as a cue to discriminate friction (Spearman's coefficient = 0.14, $p = 0.2$) (Fig.S7C).

Normal pressure results in a force vector angle which depends on the vibration amplitude (ANOVA, F(6,1593) = 67.9, p¡0.001) (Fig.S7B). The peak force angle is on average 9.9°± 4.9°SD when the friction is high and 4.2°± 2.1°SD when the friction is low. This suggests that in high-friction cases, tangential forces induced at the interface limit the global displacement of the finger, whereas in low-friction cases, tangential forces are released and micro-slips occur. However, the force vector angle is not correlated with the participants' answers (Spearman's coefficient = 0.24, $p = 0.03$), which suggests it was not used as a cue (Fig.S7D).

#### Influence of real contact area variation

The area of real contact directly influences the frictional strength of the contact, i.e. the maximal lateral force the contact can support (12). On the other side, a smaller real area of contact causes larger divergence of skin deformation when the finger is compressed against the plate. However, the real contact area difference does not succeed to predict participants' answers since we found no significant correlation between the both metrics (Spearman's coefficient = -0.07, $p = 0.49$).

#### Surface skin strains and strain energy

Strain components are shown for a low- and a high- friction case in Fig.S8B and D respectively and movie M4 shows the temporal evolution of those strains. Median strain components for each vibration amplitude are plotted in Fig.S8E. They are all positive, suggesting a skin expansion once the contact is made both in the high- and the low-friction condition. Nonetheless, the strain amplitude is larger when friction is low. Strains along $x$ and $y$ follow the same behavior as the divergence: the growth is notable at the early stage of fingertip compression and hits an inflection point when the normal force reaches 1 N. Above 2 N, the curves flatten, likely due to saturation of the compression of the fingertip pulp (10, 13). The median strain rates were computed for each of vibration amplitudes by differentiating each strain component with respect to time. They peak at the very beginning of the normal pressing when the normal force reaches $0.37 \pm 0.7$ N (Fig.S8F). We found a strong linear correlation between the median longitudinal strains and the divergence (Pearson's coefficient = 0.78) (Fig.S8G).

The strain energy densities along the skin surface were computed using (2) and are shown for a typical trial in Fig. S9A. Total strain energy follows the same behavior as the divergence with a plateau after 2 N (Fig.S9B). The action of pressing down against the surface stores mean=$0.32 \pm 0.52$ mJ of strain energy. As for strain components, there is a strong correlation between the total strain energy and the vibration amplitude (ANOVA, $F(6,1481) = 3.2$, $p = 0.004$). The median strain energy rates for each vibration amplitude are shown in Fig. S9C. These rates peak at $1.2 \pm 0.2$ mJ/s when the normal force reaches $0.4 \pm 0.1$ N. The linear correlation between total strain energy and divergence is plotted in Fig. S9D. Its slope varies from one participant to another (slope = $3.6 \pm 2.3$ mJ) because the Young's modulus we chose for this calculation does not fit for every participant of the study.

Finally, median longitudinal strain (Fig. S8H) and resulting strain energy differences (Fig. S9E) are not correlated with participants' answers. However, we found a weak correlation between the strain rate and the probability of answering that the comparison is "more slippery" (ANOVA, $F(5,70) = 2.12$, $p = 0.023$), suggesting that a sufficient deformation speed is required to enable subjects to sense frictional differences (14).

#### Strains at the depth of the mechanoreceptors

To estimate the stress inside the tissues, the skin can be modeled as a viscoelastic semi-infinite half plane (9). In this context, the spatiotemporal stimulation at the surface is spatially filtered by continuum mechanics, which diffuses stresses $\sigma(y,t)$ deeper in the soft tissues, where the mechanoreceptors are located. These stresses change consequently the local strains, following a linear first-order viscoelastic relaxation, resulting in a temporal filtering of the original stimulation.

To compute the strain to which the mechanoreceptors are sensitive to (15), the model first calculates the stress using Boussinesq and Cerruti equation (16). This model considers the skin as a semi-infinite homogeneous elastic medium on which a localized normal pressure $p(y,t)$ and tangential traction $q(y,t)$ are applied. The equation (11) leads to the shear and orthogonal normal stresses as a function of their position $y$ and depth $z$ as follows:

$$\begin{aligned} \sigma_y &= -\frac{2\,z}{\pi}\int_S \frac{p(s,t)(y-s)^2 ds}{((y-s)^2+z^2)^2} - \frac{2}{\pi}\int_S \frac{q(s,t)(y-s)^3 ds}{((y-s)^2+z^2)^2} \\ \sigma_z &= -\frac{2\,z^3}{\pi}\int_S \frac{p(s,t) ds}{((y-s)^2+z^2)^2} - \frac{2\,z^2}{\pi}\int_S \frac{q(s,t)(y-s) ds}{((y-s)^2+z^2)^2} \end{aligned} \tag{11}$$

Because of the mechanics, the spatial distribution of the pressure profile at the surface is diffused in the deeper layer of the skin. Consequently, the resulting stresses spread over a larger area and the high spatial frequency content are attenuated. The pressure and traction applied on the skin surface during a simple press on a high and a low friction surface were computed with the mechanical model detailed in the section Skin deformation model and plotted in Fig.S10B. The stress profile deep in the skin tissue are shown in Fig.S10C. Thus, the mechanoreceptors located 2 mm under the skin surface will be subjected to a resulting stress 20% higher in the high-friction than in the low-friction case (Fig.S10D).

The stresses induce a deformation of the body, following the viscoelastic Hooke's law. The compressive and shear strains $\epsilon$ can be expressed, in the Laplace domain, as a function of the local stresses:

$$\begin{bmatrix} \mathcal{L}(\epsilon_x) \\ \mathcal{L}(\epsilon_z) \end{bmatrix} = \frac{1}{E*}\begin{bmatrix} 1 & -\nu \\ -\nu & 1 \end{bmatrix}\begin{bmatrix} \mathcal{L}(\sigma_x) \\ \mathcal{L}(\sigma_z) \end{bmatrix} \tag{12}$$

where $\mathcal{L}$ is the Laplace transform, $\nu$ is the Poisson's coefficient and $E* = E + s\eta$ is the complex Young modulus of the skin layers, with $E = 1.1$ MPa the elastic modulus and $\eta$ is the viscosity of the skin and $s$ the Laplace operator. Time variation of the strain is computed numerically using a $4^{\text{th}}$-order Runge-Kutta solver. The viscoelastic behavior leads to a low-pass filtering of the surface pressure with a cut-off frequency set to $E/\eta = 100$ Hz. The tangential strains 2 mm below the skin surface are plotted in Fig.S10E and Fig.S10F shows the time evolution of the total strain at the interface, which is again 20% higher in the high-friction case.

This spatiotemporal model based on skin viscoelasticity leads to a measure of the shear and compressive strains the mechanoreceptors are subjected to. This 20% difference between a high and a low friction case is in the same order of magnitude of the just-noticeable difference typical for somatosensory system, suggesting that signalling differences between two frictional conditions is possible.

#### Ideal Observer Analysis

To test the contribution of each variable as a predictor of friction differentiation ability, we computed the performance of an ideal observer. The following variables were tested: divergence, force angle, global displacement, strain rate, strain energy density, force rate, and real contact area. Since global displacement and force angle were undesired in the experiment, the other metrics were set to NaN (not a number) for trials that present a global displacement higher than 0.2 mm, in order to evaluate the contribution of these variables when no other cues were available. The incorrect trials (i.e. when participants answered that the reference was more slippery) were first separated from the correct trials, (i.e. when they answer the comparison was more slippery). Each of the variables was normalized according to the 0.9 quantile and grouped in bins of 0.05. We counted the correct/incorrect instances in each bin and fit a normal distribution to it. The sensitivity indexes ($d'$) were extracted from the means ($\mu_1$ and $\mu_2$) and standard deviations ($\sigma_1$ and $\sigma_2$) of the Gaussian distributions of the normalized variables for correct (green) and incorrect (red) trials, as represented in figure S11A.

The probability of hits is given by the proportion of correct trials for which the variable produces a response greater than a criterion, whereas the probability of false alarms is the proportion of incorrect trials for which the variable exceeds the criterion. The receiver operating characteristics (ROC) were computed from the probability of hits as a function of the probability of false alarms when the criterion ranges from 0 to 1. The ROCs are shown in figure S11B for all tested variables. The larger the area under the curve, the better the predictor. The sensitivity index $d'$ and the area under the curve (AUC) of the ROC are summarized in table S11C. The performance of the ideal observer was on a par with the performance of the participants of the psychophysics experiments. Amongst all tested variables, the divergence metric leads to the highest sensitivity index and the highest AUC, suggesting that it is the best predictor amongst the others studied. It is followed by the force angle and the global displacement, indicating that undesired minor lateral motion present in some trials also facilitated the friction discrimination task. On the contrary, the low values of $d'$ and AUC obtained for real contact area, strain rate, SED, and force rate, mean that the participants perform at chance according to those metrics, or they possibly may even interfere with correct judgement, confirming our findings that the divergence was the most relevant metric predicting participants performance. Note that in the case of an ideal observer, choosing a criterion of 13.1 µm/mm for the divergence leads to a probability of hits of 75%.

#### Individual performance

The median of all divergence difference between reference and comparison was computed for each participant. Since the distribution is bimodal (see Fig.S12A), we divided the population of subjects into two groups. The first group contains the participants with small medians of divergences and therefore stiffer skins. The second group shows higher divergences, physically meaning larger deformations. We believe that the main contributor of the difference between the two groups is a difference in skin stiffness, as softer skin deform more under similar loading.

In both cases, the divergence difference increases with the probabilities of answering comparison is "more slippery" (Fig.S12B). Nevertheless, we observed that the group with softer skin has higher probabilities for the small vibration amplitude $\alpha <= 1$ µm than the group with stiffer skin (Fig.S12C).

#### Curl of the displacement field

The curl is a vector denoted infinitesimal rotation of a vector field. In our case, the curl is directed along the z-axis and is computed as following:

$$\int_S \nabla \times \overrightarrow{u}(x,y)dS = \int_S \frac{\partial u_y}{\partial x} - \frac{\partial u_x}{\partial y} dS \overrightarrow{z} \tag{13}$$

Similar to the divergence, the curl of the deformation field is computed for each point of the apparent contact area. A typical curl distribution is shown on Fig.S13A and the median curve is plotted Fig.S13B.

More interestingly, we found that divergence and curl follow similar trend with the normal force. Both metrics are positively linearly correlated, with a Pearson's correlation coefficient of 0.7726 ($p < 0.0001$). A possible explanation of this phenomenon postulates that fingerprints align with the direction of the stimulus, as observed in (7).

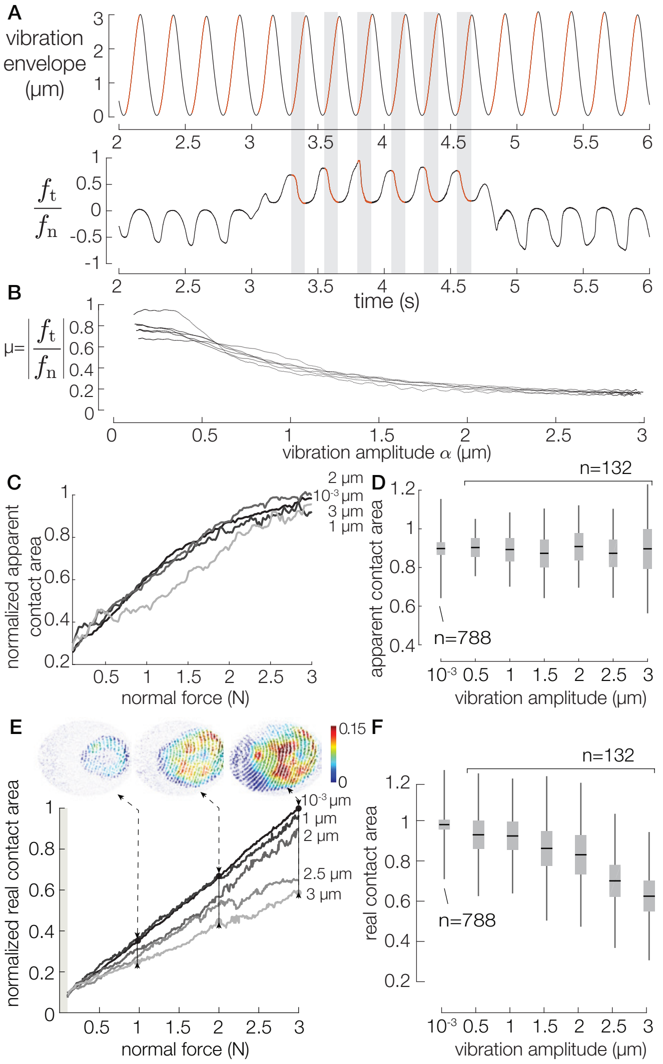


**Fig. S1.** **A**. Typical trial during the measurement of sliding friction. Subsets of time series were selected when the finger was moving from right to left and the vibration envelope as increasing. **B**. Effect of vibration amplitude on the friction coefficient. **C**. Evolution of the normalized apparent area of contact with the normal force. **D**. Median normalized apparent contact area for a normal force of 3 N. Black lines and grey boxes represent mean ± SD. **E**. Influence of the vibration amplitude on the normalized real area of contact. Images of contact area differences between the higher and lower levels of friction are shown for 1, 2, and 3 N. **F**. Median normalized real contact area for a normal force of 3 N. Black lines and gray boxes represent mean ± SD.

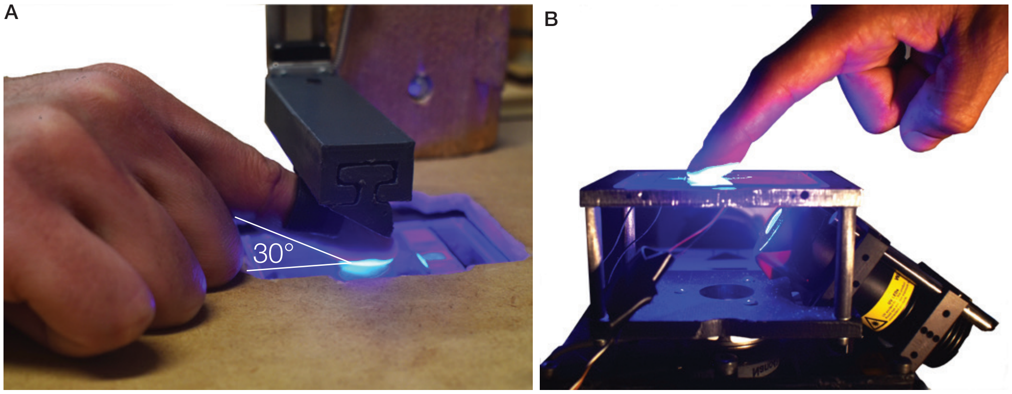


**Fig. S2.** Pictures of the experimental setup. **A**. Linear rail to maintain the angle between the finger and the surface constant equal to 30°. **B**. Friction reduction device with custom-made optical system.

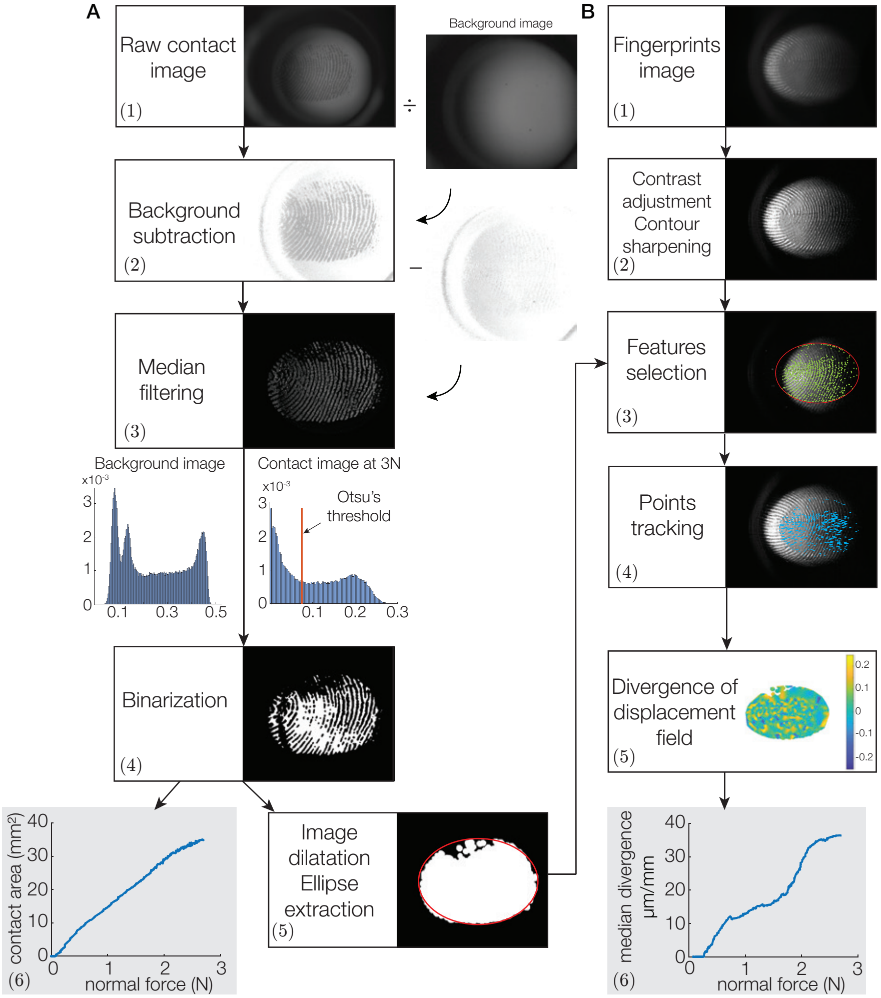


**Fig. S3.** **A**. Processing stages of the contact area: (1). Raw image of the contact. (2). Raw image normalized by the background image to correct the non-uniform lighting. (3). Contact image subtracted from the image at the first instant of contact and filtered with a median filter of radius 5. (4). Binarized image with Otsu's threshold obtained from the histogram of the filtered image of contact. (5). Opened image (dilated with circle of radius 8) using grey-scale mathematical morphological transform of Matlab and ellipse extraction. (6). Contact area in $mm^2$ as a function of the normal force. **B**. Stages of processing of the divergence: (1). Raw image of the full finger. (2). Processed image of the full finger with contrast adjustment and contour sharpening. (3). Optimal features selection at the first instant of contrast. (4). Features tracking using Lucas & Kanade algorithm. (5). Computation and interpolation of the divergence in the apparent contact area. (6). Median of divergence as a function of the normal force.

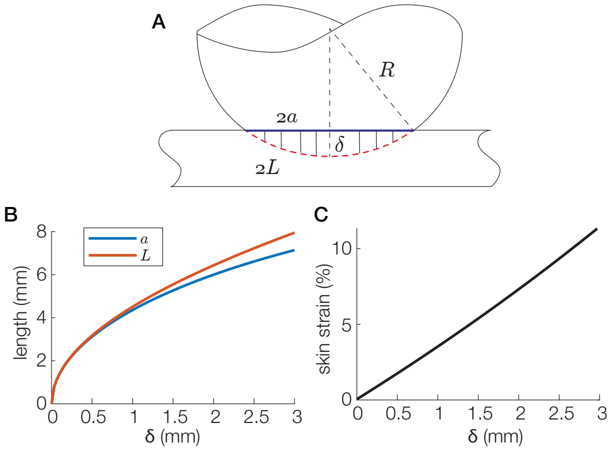


**Fig. S4.** **A**. Geometrical model of the finger, $2L$ is the length of the arc of the skin and $2a$ is the length of the contact area. **B**. Length $L$ and $a$ as a function of the normal indentation $\delta$. **C**. Skin strain in % as a function of $\delta$.

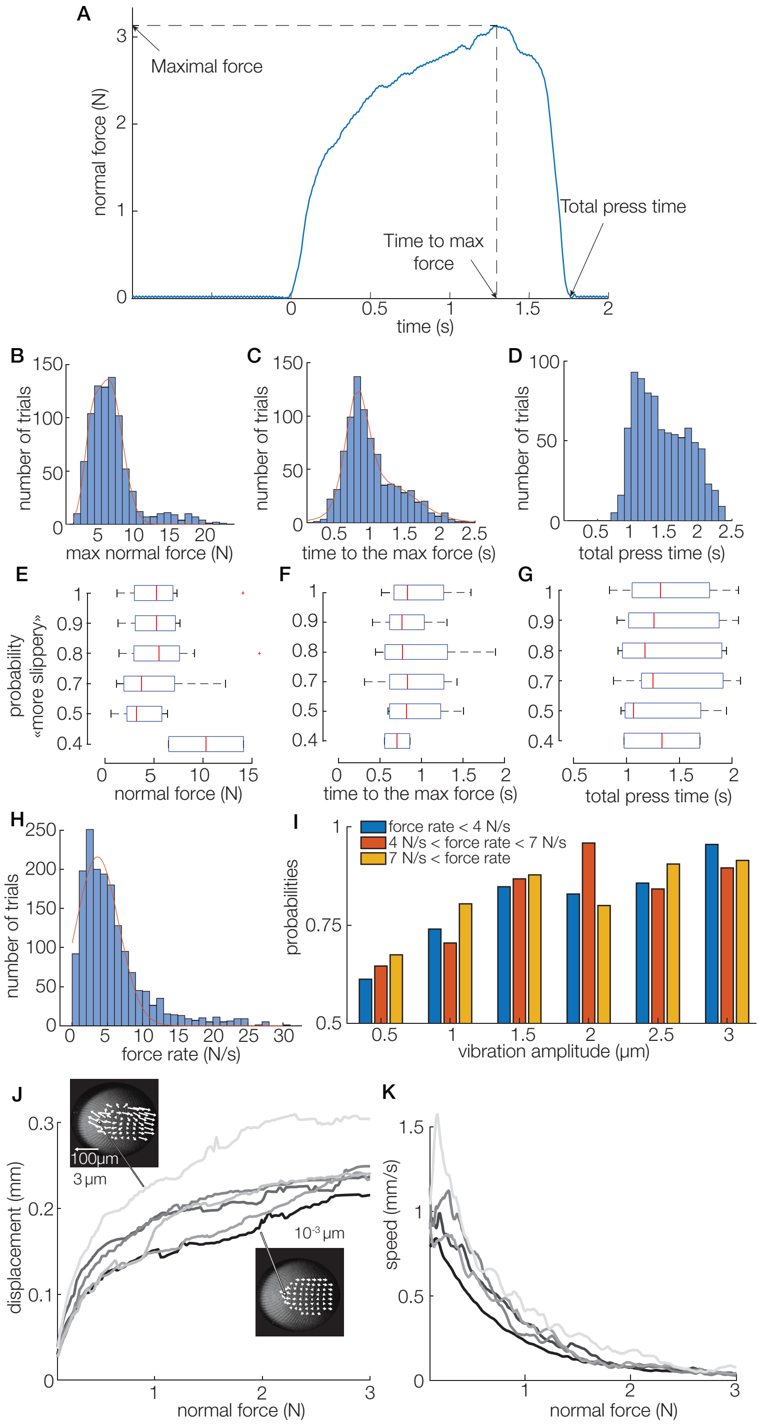


**Fig. S5.** **A**. Typical normal force time series for one trial. Distribution of normal force **B**, time to reach the maximal force **C** and total duration **D** for all trials. **E F G**. Influence of these metrics on the probabilities to answer comparison is "more slippery". **H.** Distribution of maximum force rate for all trials. **I.** Influence of the maximum force rate on the probability to answer comparison is "more slippery" for each vibration amplitude. **J.** Median maximal displacement for each vibration amplitude. **K.** Median speed of the asperity which experienced the biggest displacement along the normal force.

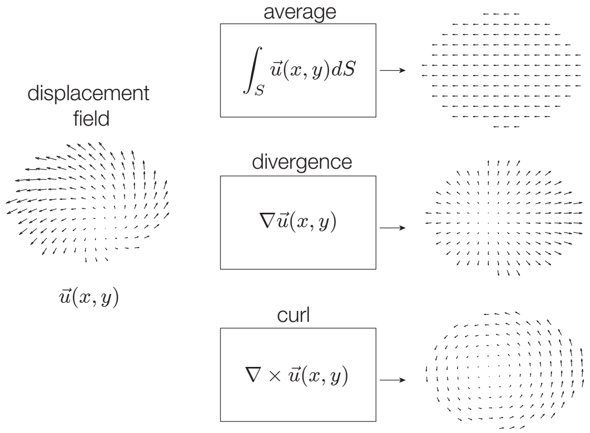


**Fig. S6.** Decomposition of a displacement field into its conservative components.

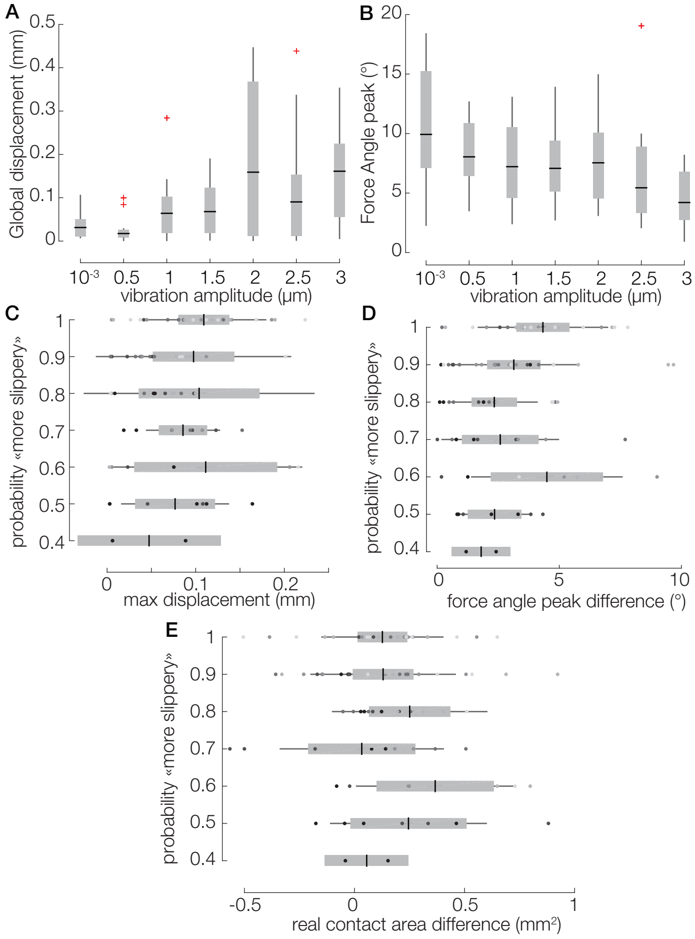


**Fig. S7.** Global displacements, given by the median value of the travelled distance by all tracked points (**A**) and force angle peaks (**B**) of all trials as a function of the vibration amplitude. **C**, **D** and **E**. The probabilities to answer comparison is "more slippery" are plotted against the median of maximal global displacements, peak force angle difference, and real contact area difference respectively. The color of the dots represents the vibration amplitude. The darker there are, the smaller the vibration amplitude.

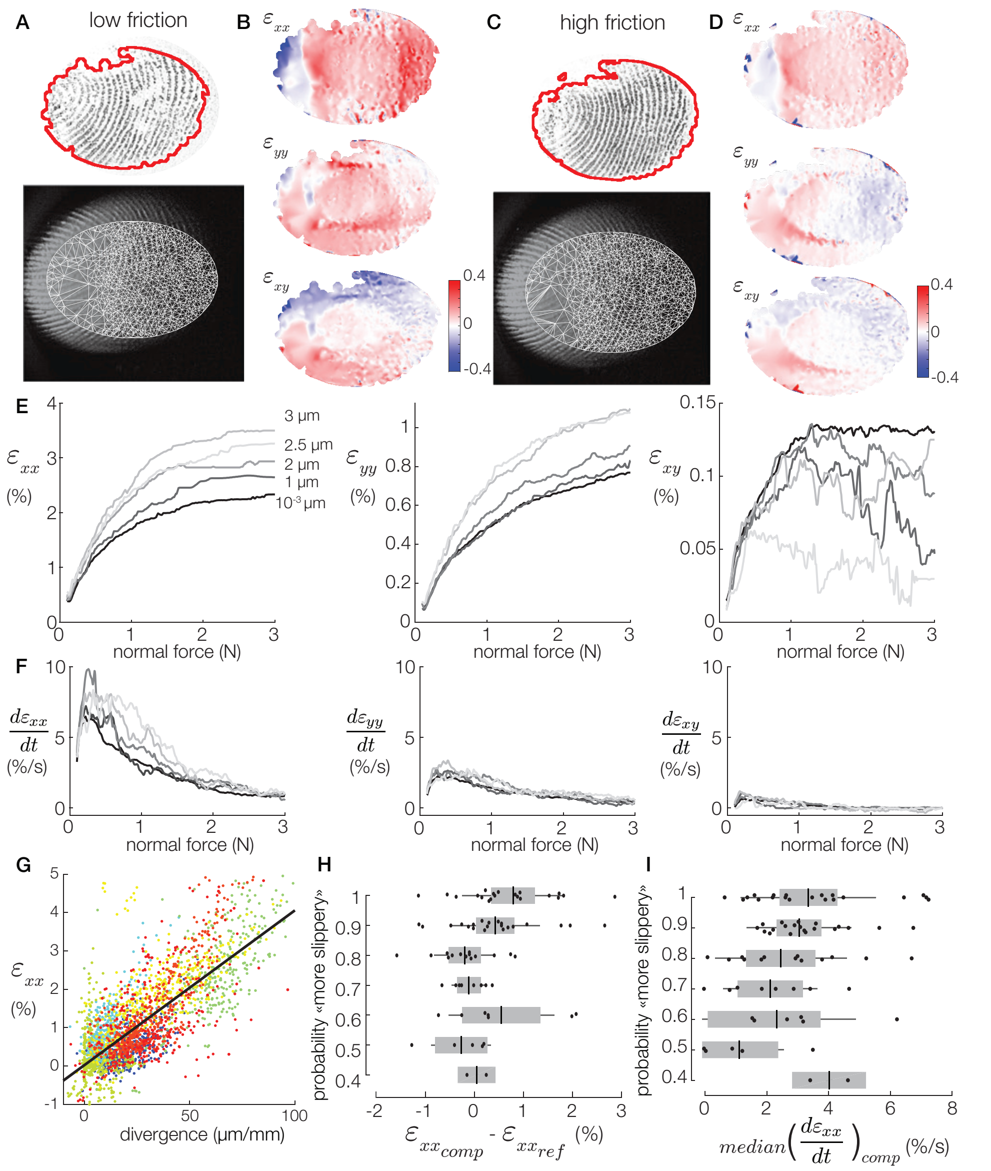


**Fig. S8.** Raw contact image and fingerprint image with the Delaunay triangulation built from the tracked points in a low friction condition (**A**) and in a high friction condition (**C**). Three strain components ($\epsilon_{xx}$, $\epsilon_{yy}$ and $\epsilon_{xy}$) represented as heatmaps for a low friction condition (**B**) and a high friction condition (**D**). **E**. Median strain components ($\epsilon_{xx}$, $\epsilon_{yy}$ and $\epsilon_{xy}$) for each vibration amplitude as a function of the normal force. **F**. Median strain components rate in %/s for each vibration amplitude. **G**. Correlation between longitudinal strain and the divergence metric ($y = 0.040x - 0.225$, $R^2 = 0.61$). Each color stands for one participant. The probabiliy to answer comparison is "more slippery" is plotted against the median of longitudinal strain difference (**H**) and the median strain rate of the comparison stimulus (**I**).

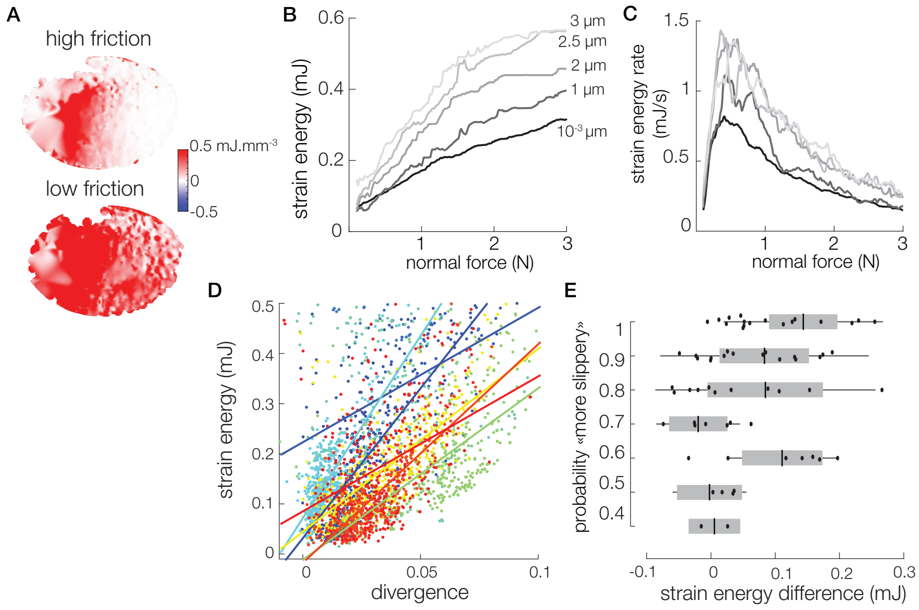


**Fig. S9.** **A**. Strain energy density in mJ.mm$^{-3}$ for a high and a low friction conditions. Total strain energy (**B**) and strain energy rate (**C**) against normal force for each vibration amplitude. **D**. The correlation between total strain energy and the divergence metric depends on the mechanical properties of participant's skin. Each color stands for one participant. **E**. The probability to answer comparison is "more slippery" are plotted against the median of strain energy difference.

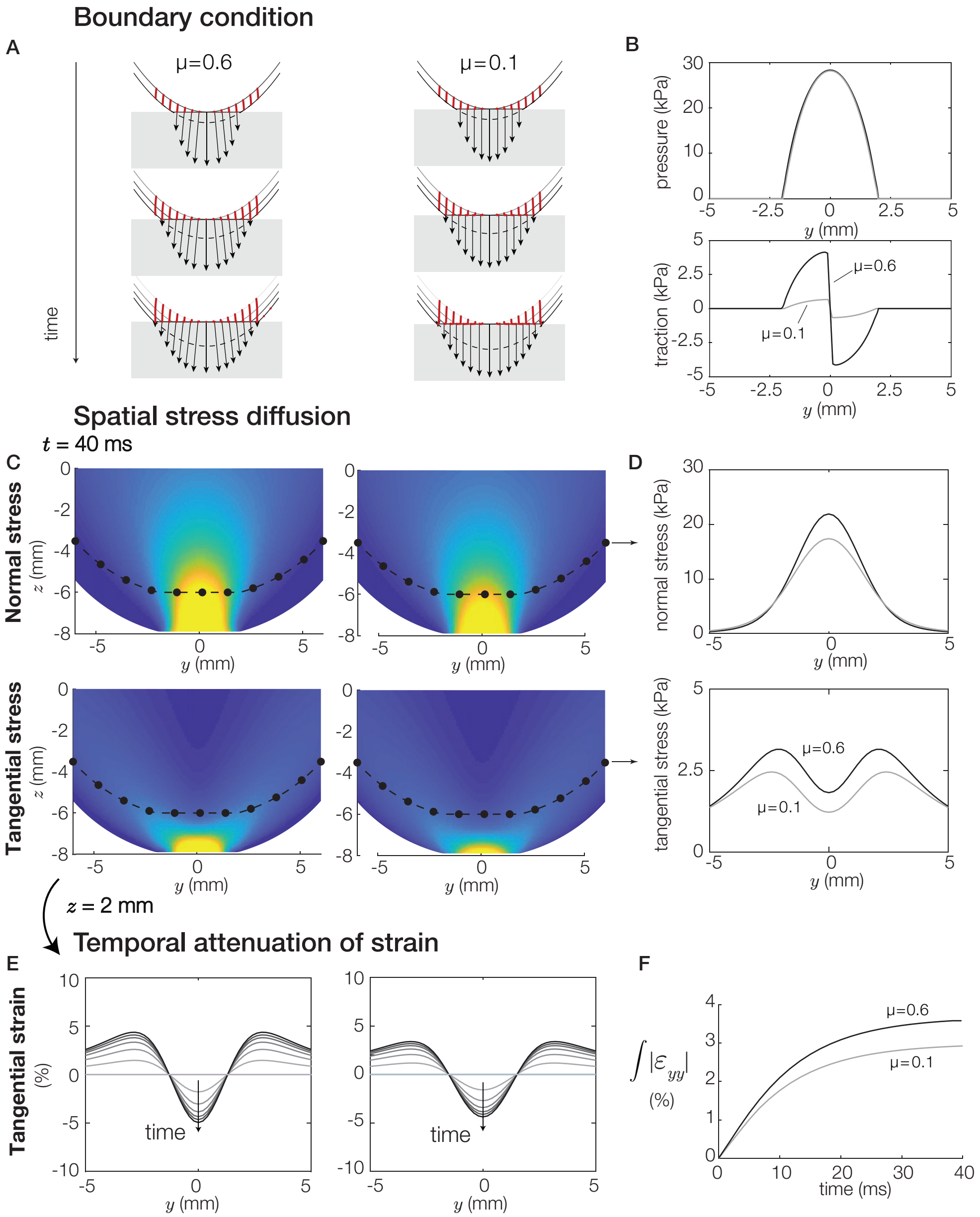


**Fig. S10.** Influence of friction on in-depth strains during a simple press. **A.** Evolution of interfacial pressure and skin deformation. **B.** Normal and tangential stresses for a high- and a low-friction condition. **C.** Spatial stress distribution inside the finger skin. The black dots correspond to the position of the mechanoreceptors, separated by 1.2 mm and 2 mm below the skin surface. **D.** Normal and tangential stresses at the mechanoreceptors' depth. **E.** Temporal attenuation of the strains 2 mm below the skin surface. We can see a dilatation of the central part and a compression aside. **F.** The internal layer of the skin is almost 20% more compressed in the high-friction case.

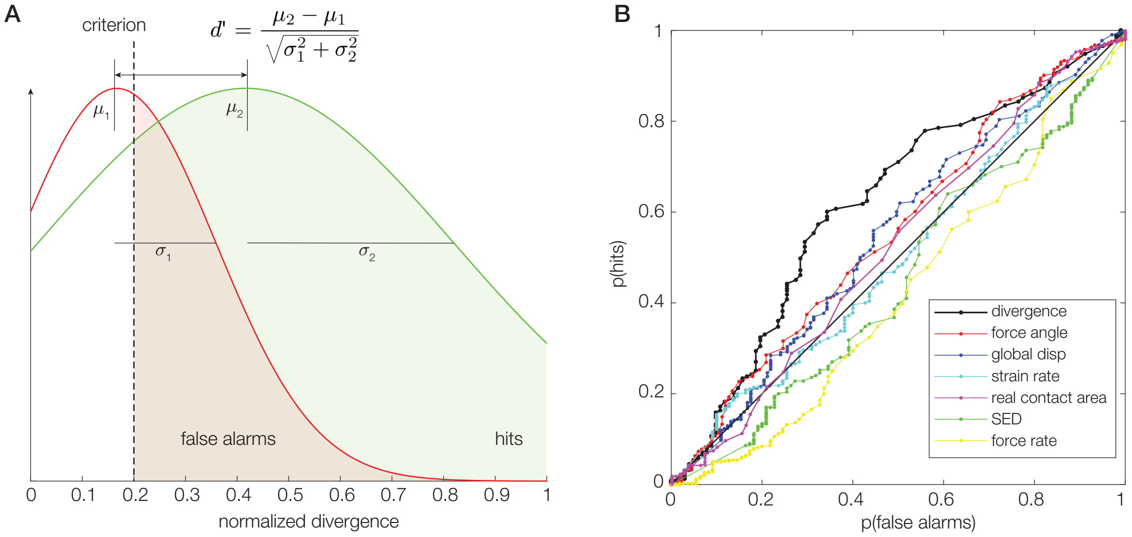


**C**

| Metric | $d'$ | AUC |
|---|---|---|
| **Divergence** | 0.47 | 0.61 |
| **Force Angle** | 0.16 | 0.56 |
| **Global displacement** | 0.14 | 0.55 |
| **Real contact area** | 0.05 | 0.52 |
| **Strain rate** | -0.05 | 0.50 |
| **Strain Energy Density** | -0.01 | 0.45 |
| **Force rate** | -0.01 | 0.42 |

**Fig. S11.** Ideal Observer Analysis. **A**. Fitted gaussian curve of the number of trials when participants are answering reference (red) and comparison (green) as a function of the normalized divergence. **B**. Receiver Operating Characteristics, representing the probability of answering the comparison when the metric is higher than a criterion (p(hits)) as a function of the probability of answering the reference when the metric is higher than a criterion (p(false alarms)). **C**. The table gathers the sensitivity index $d'$ and the area under the ROC curve (AUC) for each metric listed.

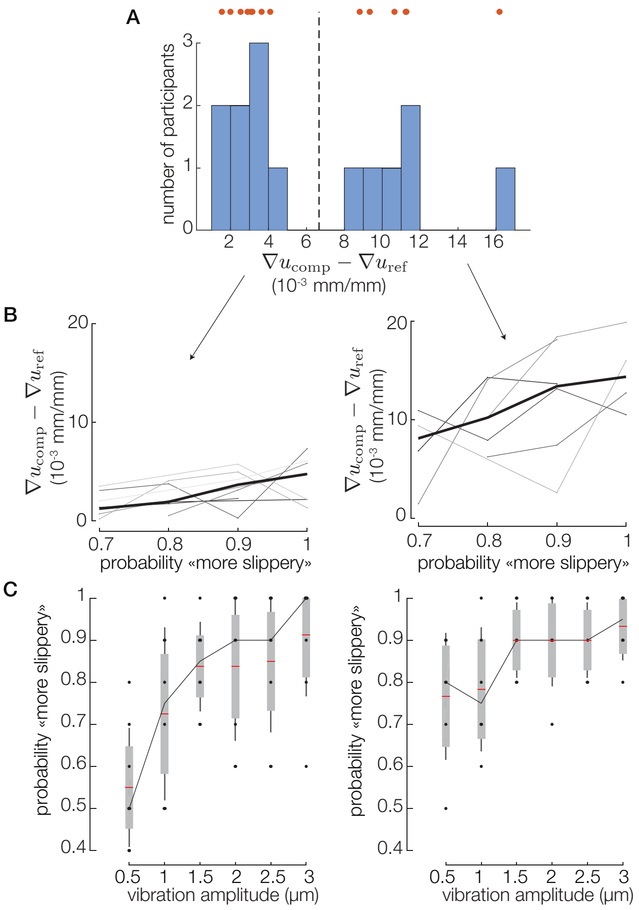


**Fig. S12.** Individual performance in friction discrimination. **A**. The histogram shows the median difference of divergences of each subject, and the dotted line divides the population of participants into 2 groups. **B**. The one on the left has a stiffer skin and experiences small deformations. The group on the right has a softer skin, experiencing more deformation. **C**. The group with softer skin has higher probability to detect the more slippery stimulus for the small vibration amplitudes, lower than 1μm.

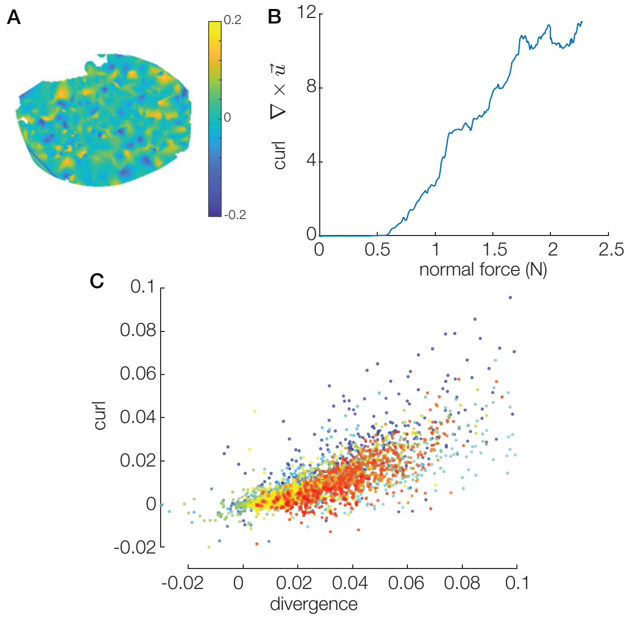


**Fig. S13.** **A**. Value of curl along the z-axis on the whole contact area. **B**. Median of curl as a function of the normal force. **C**. Correlation between median curl and median divergence. Each color stand for a participant.

#### M1. Processed images of contact area and surface features displacements for a high- and a low-friction condition.

#### M2. Experimental protocol and typical images of contact area and finger ridges obtained.

#### M3. Surface finger model deformation.

#### M4. Surface strains and strain energy density for a high- and a low-friction condition.


[1] Corentin Bernard, Sølvi Ystad, Jocelyn Monnoyer, and Michael Wiertlewski. Detection of friction-modulated textures is limited by vibrotactile sensitivity. *IEEE Transactions on Haptics*, 2020.

[8] Yuan-cheng Fung. *Biomechanics: mechanical properties of living tissues.* Springer Science & Business Media, 2013.

[9] Qi Wang and Vincent Hayward. In vivo biomechanics of the fingerpad skin under local tangential traction. *Journal of biomechanics*, 40(4):851–860, 2007.

[10] Elaine R Serina, Eric Mockensturm, CD Mote Jr, and David Rempel. A structural model of the forced compression of the fingertip pulp. *Journal of biomechanics*, 31(7):639–646, 1998.

[11] Phil R Dahl. A solid friction model. Technical report, Aerospace Corp El Segundo Ca, 1968.

[12] Nicolas Huloux, Laurence Willemet, and Michael Wiertlewski. How to measure the area of real contact of skin on glass. *IEEE Transactions on Haptics*, 2021.

[13] Dianne TV Pawluk and Robert D Howe. Dynamic contact of the human fingerpad against a flat surface. *Journal of Biomechanical Engineering*, 121(6):605–611, 1999.

[14] Benoni B Edin. Quantitative analyses of dynamic strain sensitivity in human skin mechanoreceptors. *Journal of neurophysiology*, 92(6):3233–3243, 2004.

[15] Arun P Sripati, Sliman J Bensmaia, and Kenneth O Johnson. A continuum mechanical model of mechanoreceptive afferent responses to indented spatial patterns. *Journal of neurophysiology*, 95(6):3852–3864, 2006.

[16] Kenneth Langstreth Johnson and Kenneth Langstreth Johnson. *Contact mechanics.* Cambridge university press, 1987.
